# Associations of adverse lifetime experiences with brain structure in 7003 UK Biobank participants

**DOI:** 10.1101/749077

**Authors:** Delia A. Gheorghe, Chenlu Li, John Gallacher, Sarah Bauermeister

## Abstract

**Background:** Experiences of chronic stress and trauma are major risk factors for psychiatric illness. Evidence suggests that adversity-related changes in brain structure and function accelerate this vulnerability. It is yet to be determined whether neuroendocrine effects on the brain are a result of the interference with neural development during sensitive periods or a consequence of cumulative lifetime adversity. To address this question, the present study investigated the associations between brain structure and self-reported data of childhood and adult adversity using machine learning techniques and structural equation models (SEM).

**Methods:** The UK Biobank resource was used to access Imaging Derived Phenotypes (IDPs) of grey matter and white matter tract integrity of 7003 participants, together with selected childhood and adult adversity data. Latent measures of adversity and imaging phenotypes were estimated to evaluate their associations using SEM.

**Results:** We demonstrated that increased incidence of childhood adversity events may be associated with smaller grey matter in frontal, insular, subcallosal and cerebellar regions of the brain. There were no significant associations between brain phenotypes and negative experiences during adulthood.

**Conclusions:** Using a large population cohort dataset, this study contributes to the suggestion that childhood adversity may determine grey matter reductions in brain regions, which are putatively sensitive to the neurotoxic effects of chronic stress. Furthermore, it provides novel evidence to support the “sensitive periods” model though which adversity affects the brain.

---

Arguably one of the most pressing public health problems is related to psychiatric disorders (Collins et al., 2011). An important risk factor for psychopathology are experiences of childhood adversity (CA) (Kessler et al., 2010). Furthermore, adult adversity (AA) is also linked to multiple psychiatric outcomes (Pechtel & Pizzagalli, 2011). Although the mechanisms through which adversity and chronic stress lead to these psychopathologies are not fully understood, accumulating evidence suggest a glucocorticoid effect on brain structure and function due to the dysregulation of the hypothalamic-pituitary-adrenal (HPA) axis functioning (Lupien, Juster, Raymond, & Marin, 2018). Previously, animal studies demonstrated that adverse environmental exposures during development altered the typical HPA axis response, an effect which persists throughout the life and determines structural changes in the brain (Francis et al., 1996; Meaney et al., 1991). In humans, such programming was shown to affect the morphology of the developing brain (McCrory, De Brito, & Viding, 2011), to accelerate the vulnerability to stress responsivity and psychopathology in adulthood (Pechtel & Pizzagalli, 2011), and to contribute to the neuropathology of cognitive decline in later life (Csernansky et al., 2006).

To explain this mechanism, one view posits that CA and the resulting adverse glucocorticoids affect the brain during sensitive periods of development (Lupien, McEwen, Gunnar, & Heim, 2009). For example, the hippocampus matures more rapidly in the first few years of life, while the prefrontal cortex undergoes important structural development, such as dendritic branching in response to environmental stimuli during adolescence (Giedd et al., 2009). In victims of sexual abuse, reductions in hippocampal volume were observed if the abuse occurred between the ages of 3 and 5 years old, while reductions in prefrontal cortex grey matter were associated with abuse reported during ages 14-16 (Andersen et al., 2008). Indeed, a vast amount of evidence suggests that children exposed to adversity during brain development demonstrate abnormal structural changes in limbic, prefrontal, insular, cerebellar and occipital grey matter structures, as well as in white matter tracts (Hart & Rubia, 2012; McCrory et al., 2011). Early life adversity-related changes have also been reported in the absence of psychopathology (e.g. Edmiston et al., 2011) and even with mild forms of adversity, e.g. parental discord (Walsh et al., 2014).

A well-established view suggests that cumulative and prolonged exposure is responsible for the negative consequences of adversity on the brain. Chronic stress or pharmacologically induced high levels of glucocorticoids have a cumulative neurotoxic effect on brain regions high in glucocorticoid and mineralocorticoid receptors, such as the hippocampus (Frodl & O’Keane, 2013; Sapolsky, Uno, Rebert, & Finch, 1990). In accordance with a “dose-response” effect, the allostatic load theory maintains that multiple interactive biological systems respond to prolonged and cumulative wear-and-tear (McEwen, 2004), providing one hypothesis for the negative effect. Indeed, the severity and duration of CA has been associated with larger effects on the brain. For example, the size of the left amygdala was negatively associated with the length of deprivation in institutionalized children (Mehta et al., 2009). Furthermore, reductions in visual cortices were related to the duration of sexual abuse experienced before the age of 12 (Tomoda, Navalta, Polcari, Sadato, & Teicher, 2009).

While psychopathological outcomes may be more strongly related to childhood, compared to adult adversity (Weber et al., 2008), there is currently a paucity of studies evaluating the impact of childhood versus adult adversity on brain structure. Specifically, whether the effects on the morphology of the brain are a result of cumulative adversity throughout the lifetime or the interference with development during sensitive development periods is yet to be determined. The current cross-sectional study aims to address this question by evaluating the associations between self-reported childhood and adult adversity, and derived phenotypes of grey matter structure and white matter integrity in UK Biobank participants.

## Materials and Methods

### Participants

The data in this study are from the UK Biobank resource (http://www.ukbiobank.ac.uk), a large epidemiological cohort collecting extensive genetic, biomarker, cognitive, psychometric and lifestyle measures. 502,642 participants were successfully recruited into the UK Biobank. Participants were aged 40-69 years at the time of recruitment in 2006-2010 (Sudlow et al., 2015). In 2016 the imaging phase of the study began, with the aim to scan 100,000 cohort participants by 2022 (Miller et al., 2016). UK Biobank received ethical approval from the Research Ethics Committee (11/NW/0382). Volunteers gave informed consent for their participation. The current analysis was initially conducted on 7,003 individuals: 3,593 females, aged 45-79 (M=62.04, SD=6.99) and 3,410 males, aged 46-80 (M=62.95, SD=7.27). Selection of the final sample and further details of demographics are provided in supplemental materials (Figure S1, Tables S1, S2). None of the scanned participants in the final sample had chosen to withdraw their data.

### Adverse events in childhood and adulthood

An online mental health questionnaire was administered as a follow-up self-assessment (2016) wherein selected items on traumatic events occurring in childhood or adult life were administered. CA items were based on the short version of the Childhood Trauma Questionnaire (CTS-5) (Glaesmer et al., 2013) and AA was evaluated using five bespoke questions (Khalifeh, Oram, Trevillion, Johnson, & Howard, 2015). Only items with correlation coefficients > .3 with at least one other item were retained to construct the latent models for the CA and AA questionnaires, respectively (Spinhoven et al., 2014) (Tables S3, S4). Table 1 shows the items included for this analysis. We also considered other variables, which are indicative of rearing conditions or early life factors (EL) (Allin et al., 2004; Mund, Louwen, Klingelhoefer, & Gerber, 2013; Oddy et al., 2003): birth weight, comparative body and height size at age 10, maternal smoking around the time of birth and having been breastfed as a baby (Table S2).

**Table 1.**
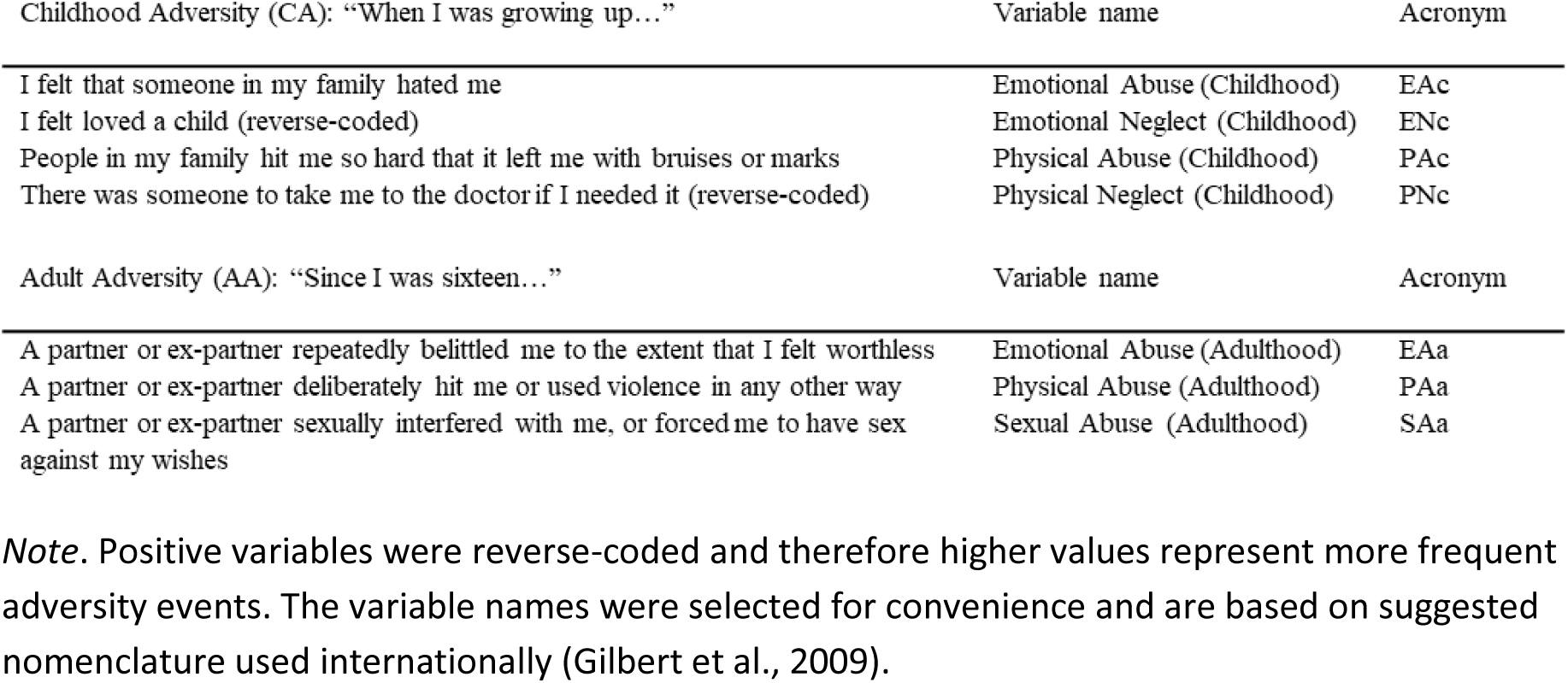

### Brain imaging

Brain images were acquired on a Siemens Skyra 3T scanner with a 32-channel RF receive head coil (Siemens Medical Solutions, Germany). Acquisition of brain imaging data and pre-processing were performed by UK Biobank. Scanning was conducted using identical protocols. In the current analytical sample 96.3% of participants (n = 6,747) were scanned in Cheadle Manchester, and the remaining (n = 256) in Newcastle. Complete details of the scanning protocol and analysis pipeline are available on the UK Biobank website (http://biobank.ndph.ox.ac.uk/showcase/label.cgi?id=100). The current study used the pre-processed derived imaging data (IDPs) (Alfaro-Almagro et al., 2018). Among the structural images, we included the total white and grey matter volumes (WM, GM), together with 139 grey matter IDPs. Of the 139 IDPs, 65 were parcelled between hemispheres (total 130). Microstructural tract integrity was investigated by looking at the diffusion-tensor-imaging (DTI) outputs: fractional anisotropy (FA) and mean diffusivity (MD). Each output mapped 27 white matter tracts (12 bilateral).

### Analysis strategy

Analyses were conducted using R V3.5.3 (R Development Core Team, 2010) and STATA V15.1 (StataCorp, College Station, Texas). All IDPs were first normalized using the head size scaling factor multiplied by each imaging variable and all bilateral IDPs were averaged. After averaging, the analysis included 15 variables for FA/MD and 74 grey matter IDPs. Observations were excluded if IDPs > ±3SD from the sample mean. Subsequently, IDPs were adjusted for age, sex, handedness, ethnicity and education, using the residual adjustment method (Ritchie et al., 2018). A principal component analysis (PCA) was used to obtain one latent measure for each type of tract measurement (global Fractional Anisotropy: gFA / global Mean Diffusivity: gMD). For grey matter IDPs, the Least Absolute Shrinkage and Selection Operator (LASSO) (Tibshirani, 1996) was used to select the IDPs which predicted either of the CA / AA items (Table S11). After identifying non-redundant features, separate PCAs were conducted on frontal, temporal, parietal, occipital, cerebellar and basal ganglia IDPs respectively (Table S12 for retained IDPs with same directionality). The Harvard-Oxford and the Diedrichsen (cerebellum) atlases were used to group IDPs (Diedrichsen, Balsters, Flavell, Cussans, & Ramnani, 2009; Smith et al., 2004). To determine the number of factors, Kaiser’s criterion of 1 and scree plots were initially explored. Subsequently, parallel analysis was used, and factors were retained if the adjusted eigenvalues were both > 1 and the randomly generated eigenvalues (Dinno, 2009; Horn, 1965). Where more than one component was identified, orthogonal (varimax) rotation was used (Costello & Osborne, 2005). Finally, SEM constructed latent measurement models of CA, AA, and EL with the Satorra-Bentler robust estimator (Satorra & Bentler, 2001) to separately predict the brain structure outcomes.

## Results

### Latent factors of white matter microstructure

White matter tracts correlated positively across the brain (FA *r* range: 0.53–0.97; MD *r* range: 0.53–0.98; all *p* < .001), suggesting consistent directionality (Tables S9, S10). A single un-rotated factor explained 79.37% (FA) and 83.23% (MD) of the total integrity variance (Figure 1). All tracts demonstrated large positive loadings for each of the two measurement types, and therefore no coefficients were supressed (Table 2).

**Table 2.**

|  | **Fractional Anisotropy** | **Mean Diffusivity** |
| --- | --- | --- |
| **Sensory projections** |  |  |
| Acoustic radiation | 0.91 | 0.92 |
| Corticospinal tract | 0.91 | 0.95 |
| Middle cerebellar peduncle | 0.78 | 0.68 |
| Medial lemniscus | 0.80 | 0.87 |
| **Thalamic projections** |  |  |
| Anterior thalamic radiation | 0.94 | 0.96 |
| Posterior thalamic radiation | 0.94 | 0.93 |
| Superior thalamic radiation | 0.92 | 0.95 |
| **Callosal Fibres** |  |  |
| Forceps major | 0.89 | 0.81 |
| Forceps minor | 0.94 | 0.93 |
| **Association Fibres** |  |  |
| Cingulate gyrus part of cingulum | 0.83 | 0.95 |
| Parahippocampal part of cingulum | 0.67 | 0.83 |
| Inferior fronto-occipital fasciculus | 0.97 | 0.98 |
| Inferior longitudinal fasciculus | 0.97 | 0.97 |
| Superior longitudinal fasciculus | 0.95 | 0.97 |
| Uncinate fasciculus | 0.89 | 0.93 |
| **Explained variance** | 79.37% | 83.23% |
| **Mean between-tract r** | 0.77 | 0.81 |
| **Overall Kaiser-Meyer-Olkin** | 0.97 | 0.97 |

*Note.* Tract loadings for each white matter measurement, explained variance for the single unrotated principal component, average correlation magnitude (Person's $r$ ) across all white matter tracts within each measurement type

**Figure 1.**
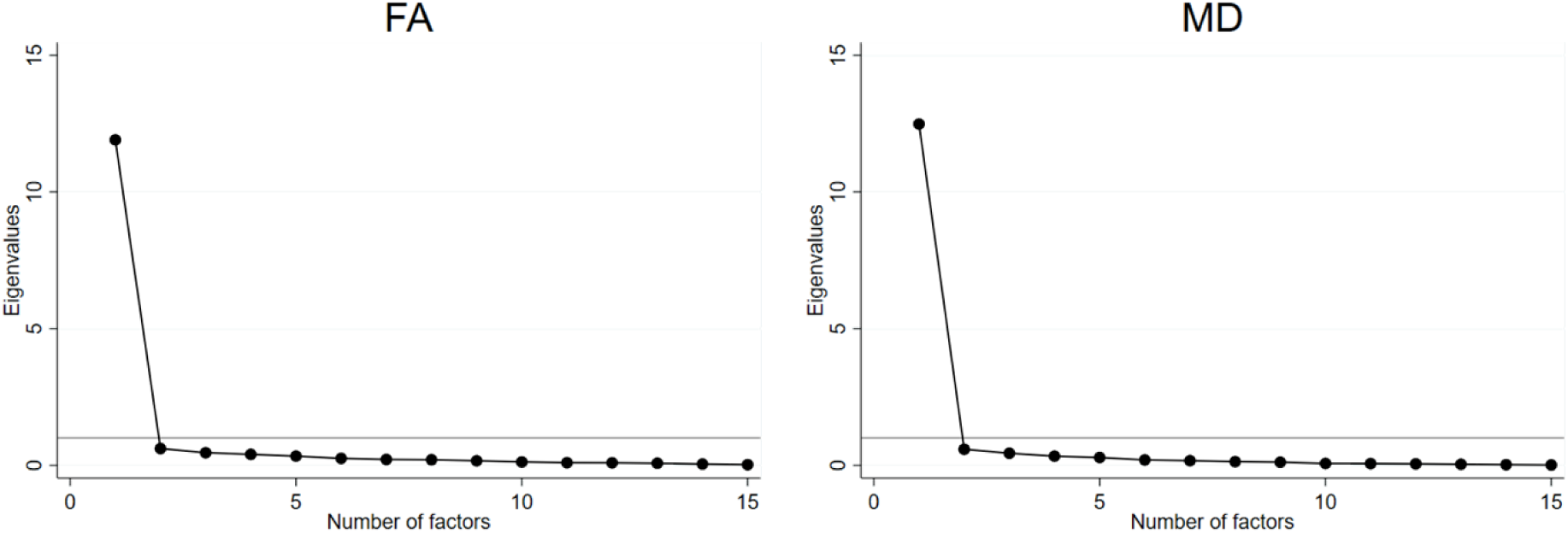
Scree plots for the PCA based on the FA and MD white matter integrity tracts. In both cases, the first factors were associated with large eigenvalues (11.91 – FA; 12.49 – MD), indicating a single latent factor.

### Latent factors of grey matter structure

LASSO identified 52 features using 10-fold cross-validation. With the exception of PNc, each item identified non-zero coefficients (Table S11). To construct the latent factors, selected variables were grouped according to their parent categories: cerebellum; basal ganglia; frontal; temporal; parietal; occipital lobes (Table 3). PCAs were conducted on each group after excluding four variables with opposite association directions: middle temporal gyrus; anterior part; middle temporal gyrus; temporo-occipital area; temporo-occipital fusiform cortex; vermis Crus I. Within each group, IDPs showed significant positive correlations, *r* range cerebellum: 0.10-0.78; *r* range basal ganglia: 0.14-0.61; *r* range frontal lobe: 0.09-0.38; *r* range temporal lobe: 0.05-0.53; *r* range parietal lobe: 0.06-0.48; *r* range occipital lobe: 0.05-0.52; all p < .01 (Table S12). One single un-rotated factor was retained for the global measure of the basal ganglia (gBG, 52% variance); cerebellum (gCBL, 47% variance); frontal lobe (gFL, 30% variance); parietal lobe (gPL, 33%) and occipital lobe (gOL, 38% variance) (Figure 2). In the temporal lobe a rotated two-factor solution explained 54% of the total structural variance. None of the coefficients were supressed due to low loadings (i.e., < .3).

**Table 3.**
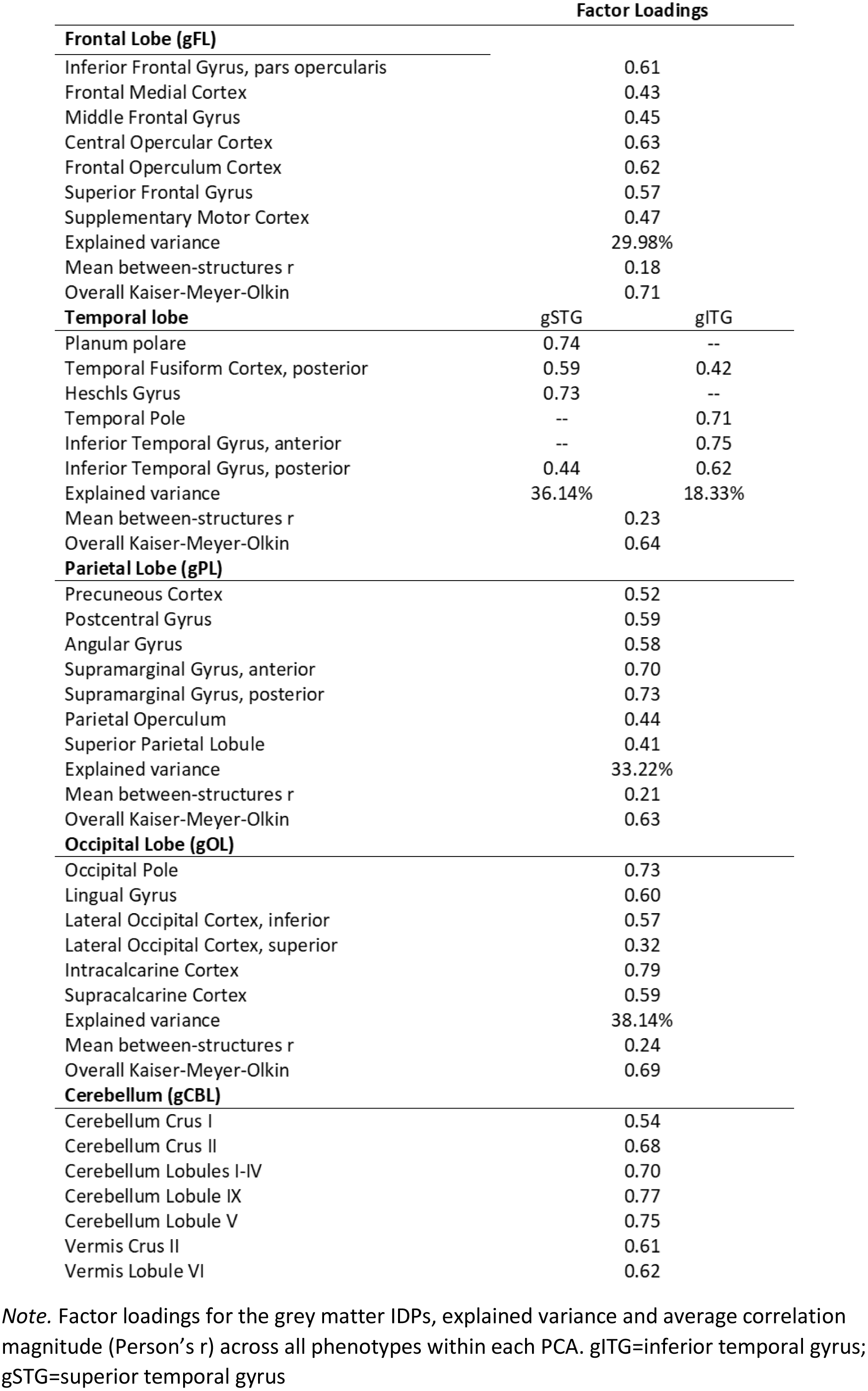

**Figure 2.**
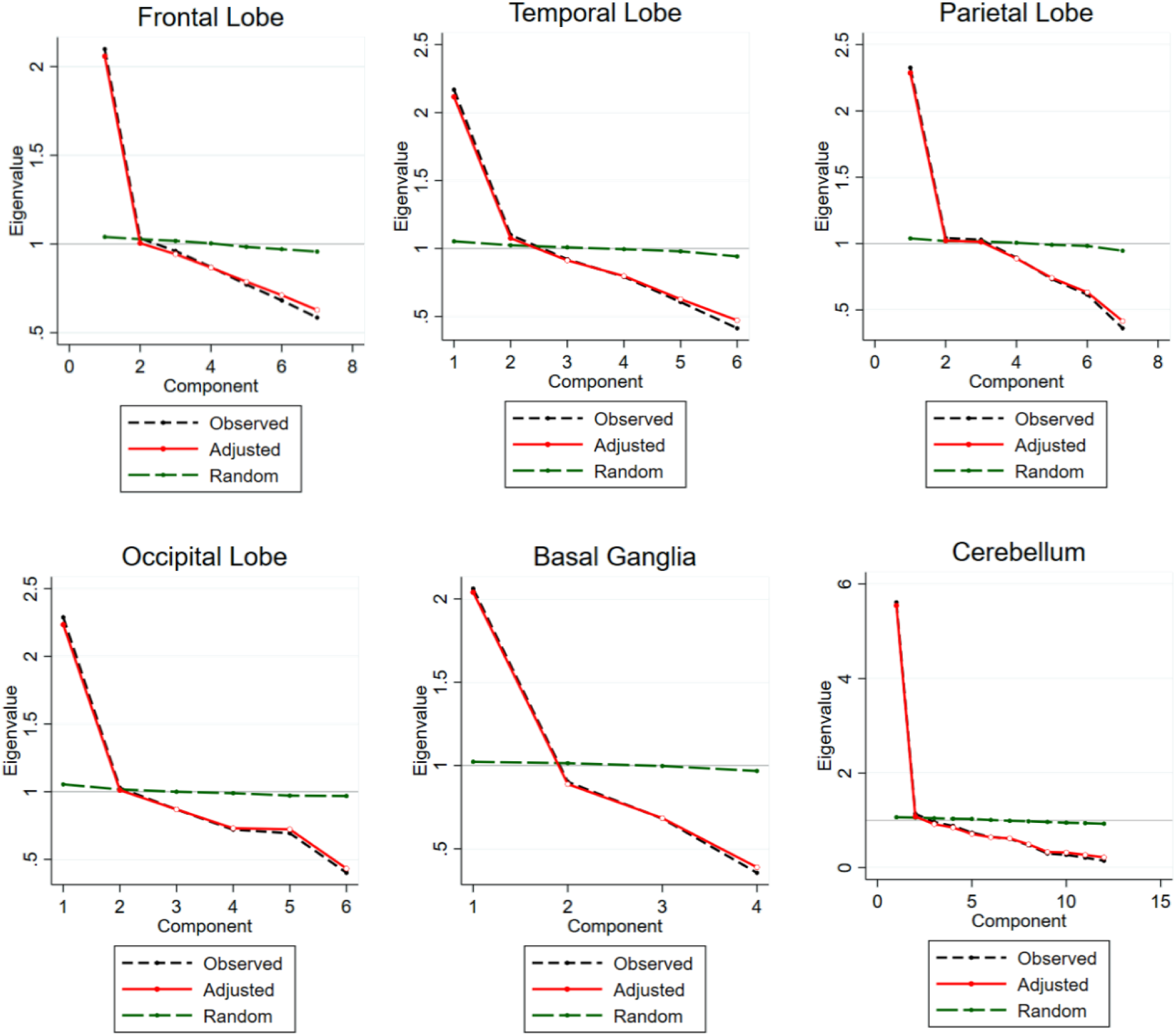
Scree plots of parallel analysis for each PCA based on the grey matter IDPs. The dashed black lines plot the observed eigenvalues reported by the PCA analysis. The red line represents the sampling error-adjusted eigenvalues obtained after parallel analysis. The green line depicts randomly generated eigenvalues. A factor was retained if the adjusted eigenvalue was > 1 (grey horizontal line) and if it was above the randomly generated eigenvalues. Apart from the temporal lobe, all remaining PCAs suggest a one-factor solution.

### Structural equation models of adversity, early life factors and brain measures

Latent brain measures and selected individual IDPs were evaluated in separate structural models against CA, AA and EL (Figure 3). Due to missingness in the EL factor variables (0.29-32.64%), the analysis was conducted using list-wise deletion (resulting N = 4,144). All structural models showed good fit (Hooper, Coughlan, & Mullen, 2008) (CFI = 0.98; TLI = 0.97-0.98; RMSEA = 0.02; SRMR = 0.02-0.03) (Table S13). Increased incidence of CA was associated with smaller grey matter in the frontal, cerebellar, subcallosal and insular regions (β=-0.04:-0.05; p<.05). AA was not significantly associated with the brain measures (β=-0.03:0.04), and there were no indirect effects on structure from CA through AA or from EL through CA (β=-0.01:0.02). No significant effects were identified between CA/AA and white matter measures (β=-0.01:0.03). Associations are summarized in Tables 4 and 5. For additional information on the development of the adversity and EL measurement models see Tables S5-8, Figure S2.

**Table 4.**
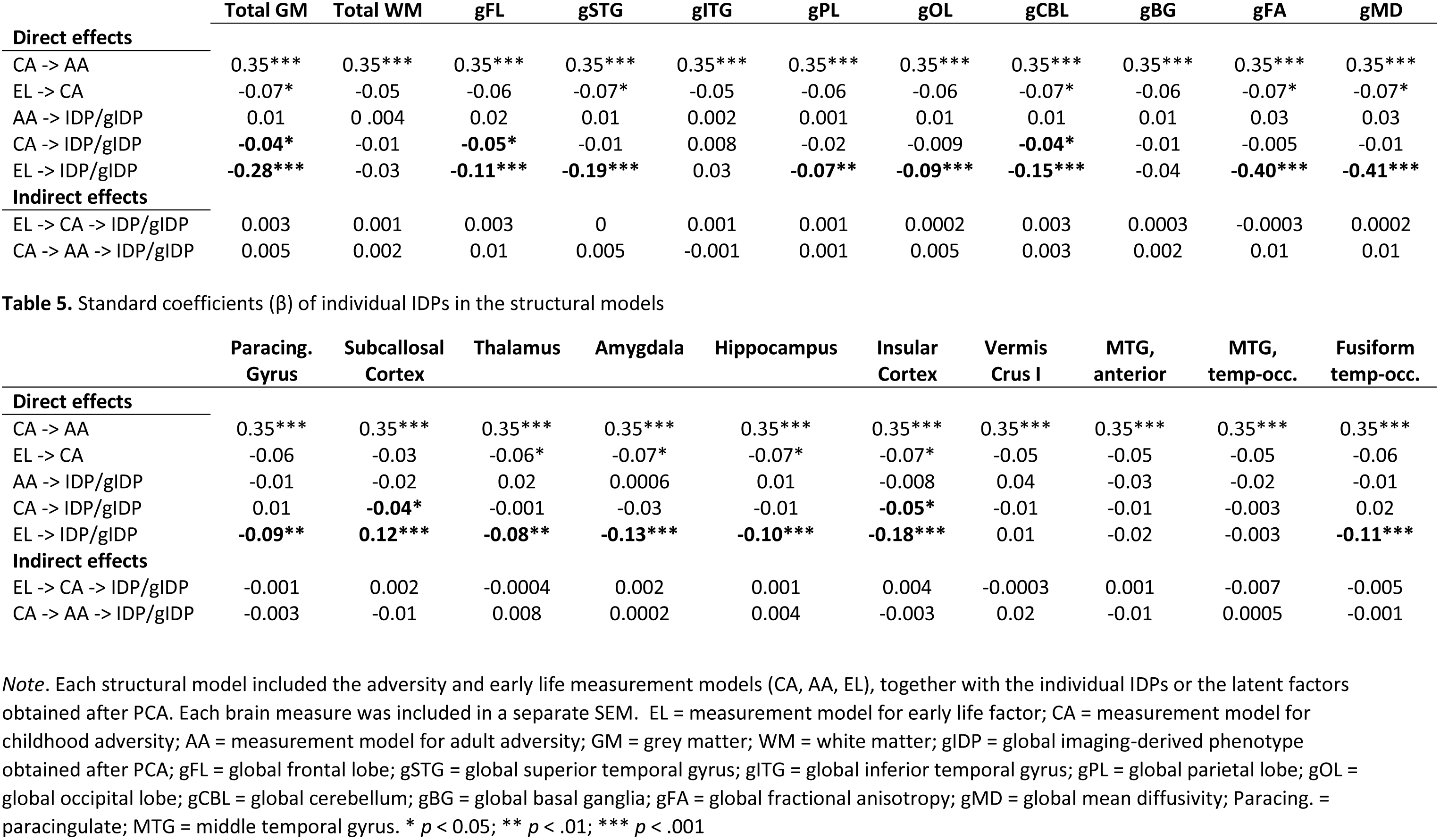
Standard coefficients (β) of the latent brain measures, total grey and white matter in the structural models

**Figure 3.**
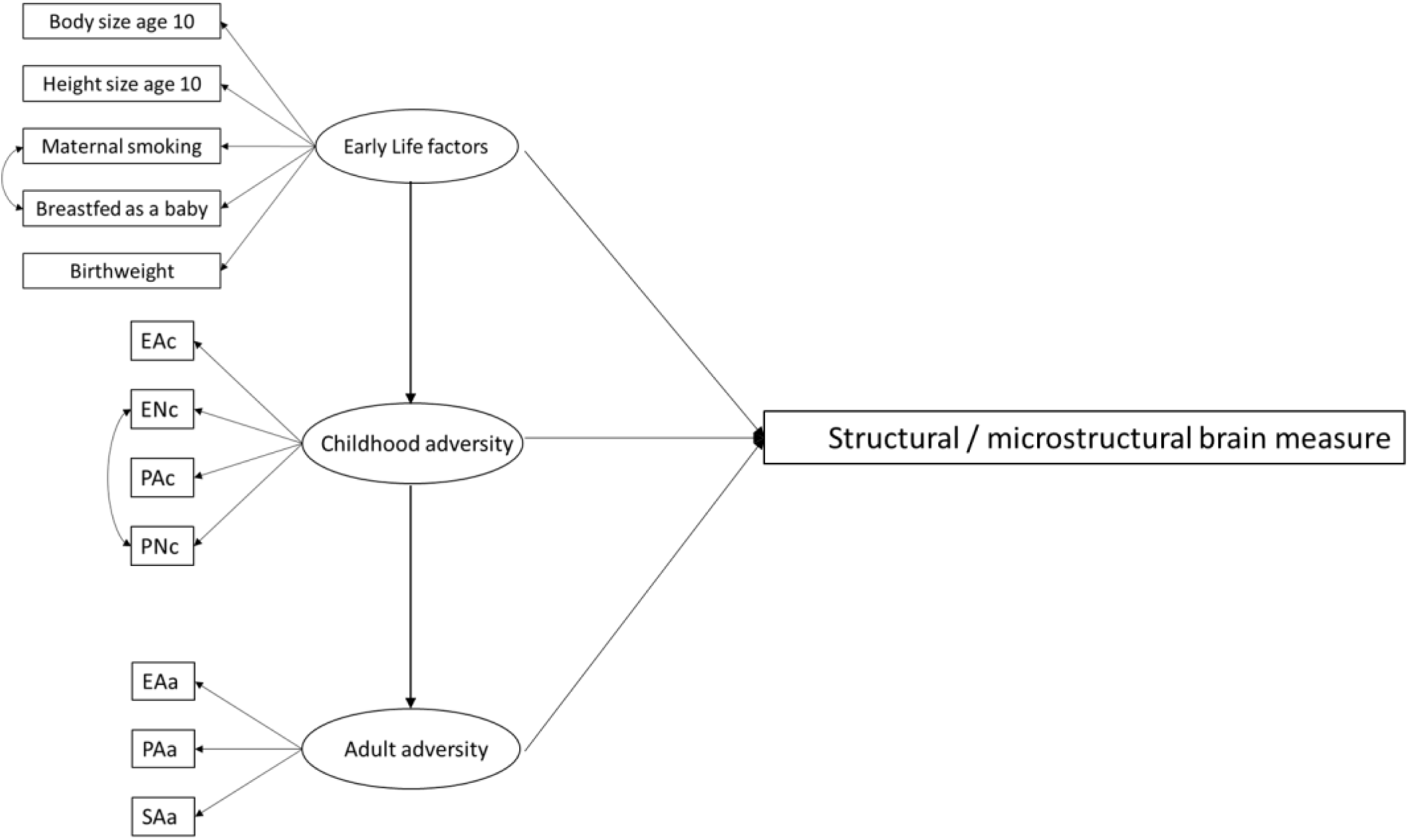
Structural equation models. In separate models, the structural/microstructural brain measure box included: total grey matter, total white matter, global factors of frontal, parietal, occipital lobes, superior and inferior temporal gyri, and microstructural integrity (fractional anisotropy / mean diffusivity). In addition, separate IDPs were also evaluated: paracingulate gyrus, subcallosal cortex, thalamus, amygdala, hippocampus, insular cortex, vermis crus I, middle temporal gyrus (anterior, temporo-occipital), temporo-occipital fusiform cortex.

## Discussion

The current study investigated associations between brain phenotypes and experiences of adversity in childhood and adulthood. The aim of this analysis was to evaluate whether the phenotypic changes are linked to the timing of adversity events. We found that smaller total, frontal, insular, cerebellar and subcallosal grey matter are associated with increased adversity experienced during childhood, but not adulthood. In addition, although there is a positive relationship between childhood and adult adversity, we found the effects of early life events on brain structure are not mediated by later life adversity. White matter variables are not associated with adversity.

Our findings are interesting considering that brain regions with modulatory roles on the stress response, may be particularly vulnerable to the effects of adversity during development (McLaughlin, Sheridan, & Lambert, 2014). Indeed, the prefrontal cortex is anatomically connected to the paraventricular nucleus of the hypothalamus (Floyd, Price, Ferry, Keay, & Bandler, 2001), which is the origin of the HPA response to stressors. Furthermore, changes in frontal areas have been reported in maltreated children (e.g. De Brito et al., 2013), as well as adults who experienced childhood adversity (e.g. Andersen et al., 2008, Edmiston et al., 2011). Similarly, the subcallosal cortex has strong anatomical connections with the medial prefrontal cortex and responds preferentially to negatively-valenced emotional content (Laxton et al., 2013). Reduced grey matter and poor functional connectivity with fear-processing areas have been reported in this region, in response to childhood maltreatment (McLaughlin et al., 2014). Further, the insular cortex, which has been involved in understanding the feeling of pain (Singer et al., 2004), showed grey matter reductions in children exposed to physical abuse (Edmiston et al., 2011). Finally, we also found that smaller cerebellar grey matter volume was associated with childhood adversity. Accumulating evidence from lesion, imaging and brain stimulation studies have demonstrated that the cerebellum supports emotional processing (e.g. Schmahmann & Sherman, 1998; Schutter & van Honk, 2006; Stoodley & Schmahmann, 2009). Furthermore, consistent reductions in cerebellar volumes have been reported in children exposed to severe, as well as mild childhood adversity (e.g. De Bellis & Kuchibhatla, 2006; Walsh et al., 2014). Importantly, the cerebellum also has anatomical connections with the HPA system via mono-synaptic projections to the paraventricular nucleus of the hypothalamus (Schutter, 2012). Considered together, adverse structural changes in frontal lobe, adjacent limbic and paralimbic structures, as well as cerebellar regions, may represent important phenotypes of early adversity.

We did not find significant associations between childhood adversity and white matter integrity. Children who have experienced severe neglect show more diffuse organisation in prefrontal white matter microstructures (Hanson et al., 2013) and institutionalized children who faced severe deprivation demonstrated reduced white matter tract coherence in frontal, temporal and parietal regions (Govindan, Behen, Helder, Makki, & Chugani, 2010). Furthermore, there is some consistency in reports of reduced corpus callosum volume in children and adolescents exposed to adversity (De Bellis et al., 1999; Teicher, Tomoda, & Andersen, 2006). It may be possible that more severe forms of adversity could be associated with white matter changes, which might not be characterized in the UK Biobank sample.

It is important to note that the latent measure of early life factors was negatively associated with the global measures of white matter diffusion, frontal lobe, superior temporal gyrus, parietal lobe, occipital lobe, cerebellum, paracingulate gyrus, thalamus, amygdala, hippocampus, insular cortex and fusiform gyrus. Indeed, maternal smoking, lower birthweight and general smaller morphology as a child are indicative of high familial adversity settings (Hill, Lowers, Locke-Wellman, & Shen, 2000; Ouellet-Morin et al., 2008). Previous research suggested that maternal smoking during pregnancy was associated with reductions in the cerebellum and corpus callosum (Bublitz & Stroud, 2012) and lower birthweight predicted decreased grey matter structures and white matter integrity in multiple areas (Allin et al., 2004). Our results are consistent with these previous findings. However, cumulative effects of EL on the brain, mediated by exposure to adversity were not observed, suggesting that in the current sample, associations may be directly linked to childhood adversity.

Finally, it is important to acknowledge a series of limitations. First, this study does not account for genetic vulnerability. Indeed, theoretical accounts of adversity highlight the importance of the interaction between environmental stressors and genetically-mediated stress reactivity (Boyce & Ellis, 2005). Second, the study assesses adversity retrospectively. Therefore, it does not account for pre-existing differences in brain structure and white matter integrity. Longitudinal measurements of adversity and brain structure will be able to disentangle the intra-individual differences in brain structure from the neurotoxic effects of adversity. Finally, the adversity information available in UK Biobank is limited, and it was collected via self-report. Given the inherent complexity of the adversity concept, the items evaluated here may not provide a comprehensive assessment of childhood and adult adversity.

In conclusion, this cross-sectional study suggests that childhood adversity is associated with reduced grey matter in frontal, limbic and cerebellar regions. There were no significant associations between adversities experienced during adulthood and brain phenotypes, and adult adversity did not mediate the effects of early exposure on the brain. These results support the idea that adversity-related structural changes may occur when the brain undergoes early neuronal growth.

## Acknowledgements

This is a DPUK supported project with all analyses conducted on the DPUK Data Portal, constituting part 1 of DPUK Application 0144.

The Medical Research Council supports DPUK through grant MR/L023784/2

## Authors’ Contributions

DG and SB conceptualised the idea. DG analysed the data, and wrote the manuscript. CL and DG analysed the phenotypic data using ML. All authors interpreted the analyses. SB and JG edited and proofread the manuscript. All authors read and approved the final manuscript.

## Key points

- The UK Biobank large epidemiological resource includes data on adversity, trauma and brain imaging
- Childhood adversity is related to reductions in grey matter structure
- The detrimental effects of adversity are stronger if events occurred during childhood, compared to adulthood
- Retrospective assessment of adversity is a limitation

