## Supplemental materials for "Associations of adverse lifetime experiences with brain structure in 7003 UK Biobank participants"

**<u>Participants and descriptive statistics</u>**

**Figure S1.** *Selection of the analytical sample*

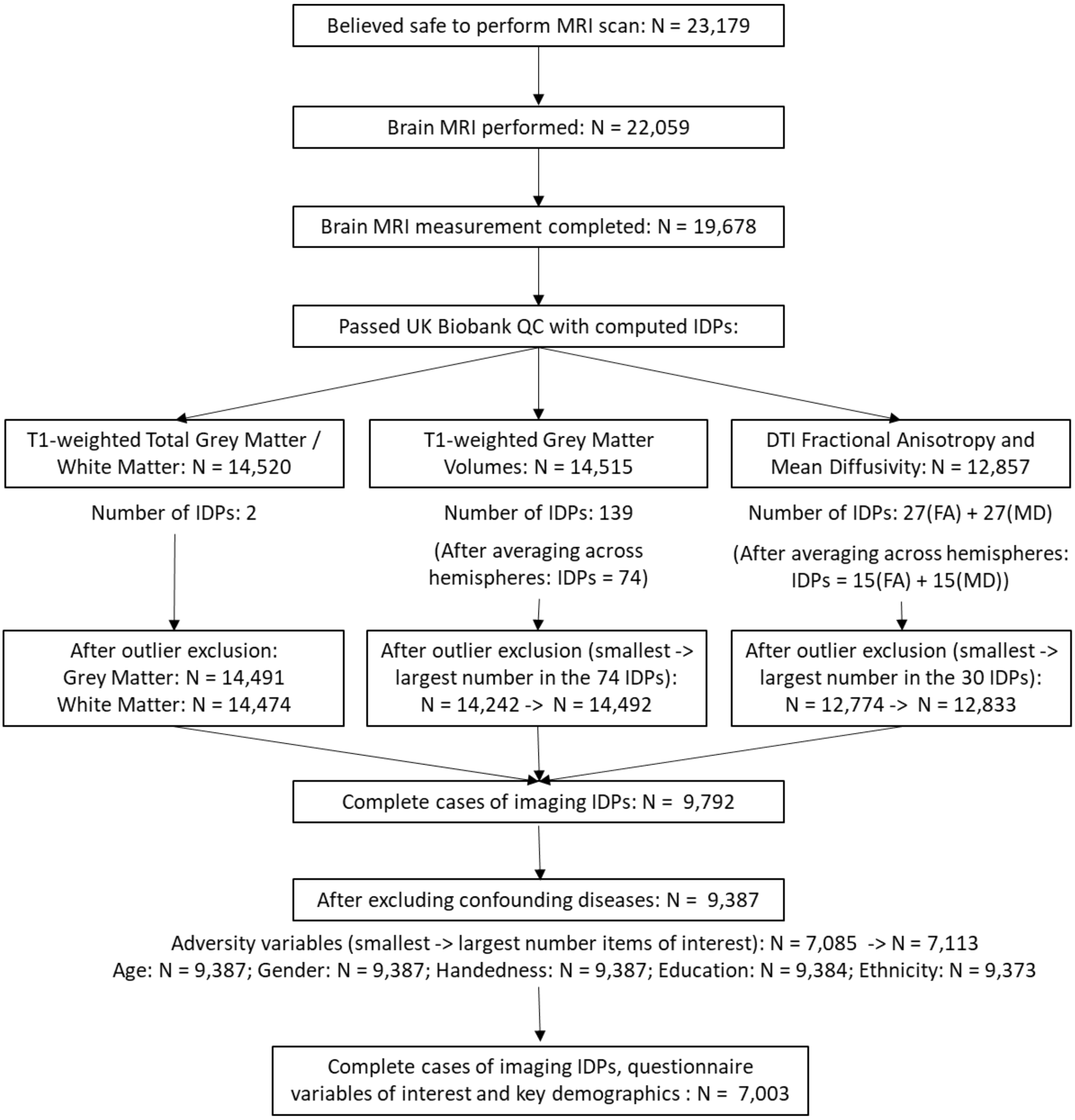


*Note.* Of the participants who volunteered to participate in the imaging, 13,179 were considered safe for scanning. Of these, the MRI scan was performed and completed for 19,678 participants. 567 of these were scanned with the early protocol phase (Phase 2), before improvements to the imaging protocol. For these scans and others which did not pass UK Biobank Quality Control (QC), Imaging Derived Phenotypes (IDPs) were not computed. After excluding outliers for each IDP of interest (averaged totals between hemispheres where applicable), and keeping observations with non-missing data on adversity variables, gender, age, handedness, education and ethnicity, the final dataset included 7,003 participants. Imaging outliers were excluded before obtaining complete cases based on the sample overlap with the adversity variables, and are therefore unbiased by these measures. Adversity items of interest were selected based on their fit in the measurement models and the inter-item correlations (Tables S3, S4, S5, S6, S8). Finally, the confounding diseases were identified based on self-reported medical conditions (http://biobank.ctsu.ox.ac.uk/crystal/field.cgi?id=20002). The following categories were included: brain haemorrhage, brain abscess, cerebral aneurysm, cerebral palsy, chronic/degenerative neurological problem, dementia/Alzheimer's/ cognitive impairment, encephalitis, head injury, infection of nervous system, ischaemic stroke, epilepsy, meningioma, meningitis, motor neuron disease, multiple sclerosis, neurological injury/trauma, benign neuroma., other neurological problems, Parkinson's disease, spina bifida, stroke, subarachnoid haemorrhage, subdural haemorrhage, transient ischaemic attack, cancer of the nervous system.

**Table S1.** *Demographics*

| | **Males** | | | **Females** | | |
|---|---|---|---|---|---|---|
| | M(SD) | Range | N | M(SD) | Range | N |
| Age [a] | 62.95 (7.27) | 46 – 80 | 3,410 | 62.04 (6.99) | 45 – 79 | 3,593 |
| Townsend [b] | -2.12 (2.61) | -6.26 – 8.71 | 3,409 | -2.06 (2.56) | -6.18 – 9.14 | 3,588 |
| | College/University degree (Yes : No) | | | | | |
| Education [c] | 1,656 : 1,754 | | 3,410 | 1,582 : 2,011 | | 3,593 |
| | Ethnic background (% white) | | | | | |
| Ethnicity [d] | 97.42% | | 3,410 | 97.89% | | 3,593 |
| | Handedness (right-handed : left-handed : ambidextrous) | | | | | |
| Handedness | 2,997 : 337 : 76 | | 3,410 | 3,240 : 298 : 55 | | 3,593 |

*Note.* [a]Age recorded at the time of the MRI scan. Where this information was missing (n = 294 of total n = 7,003) it was computed based on the date of birth and the date on which participants attended the assessment centre. Note that only month and year of birth are available and therefore, the first day of the month was considered for all these imputations.

[b]Townsend refers to the Townsend Index of neighbourhood deprivation levels (Townsend, 1987) measured at baseline.

[c]Education recorded at baseline. Coding refers to College/University degree or either of the following: A levels/AS levels or equivalent; O levels/GCSEs or equivalent; GSEs or equivalent; NVQ or HND or HNC or equivalent; other professional qualifications. Where this information was missing (n = 540 of total n = 7,003) but recorded at subsequent assessment waves, the value recorded subsequently was imputed into the education variable. The variable was dichotomized as above. Where further data was missing, this was imputed as follows: where age of completed education was > 18 and <= 18, the observation was allocated to the College/University degree group or the No College/University degree group.

[d]Ethnicity recorded at baseline. Most study participants were of white ethnicity (98% in n = 7,003). Therefore the data was dichotomised into White – Non-white, given the following categories: White (White-British, White-Irish, Any other white background); Mixed (White and Black Caribbean, White and Black African, White and Asian, Any other mixed background); Asian (Asian and Asian British,

Indian, Pakistani, Bangladeshi, Any other Asian background); Black (Black and Black, British Caribbean, African, Any other Black background); Chinese; Other ethnic group. Where this information was missing (n = 14 of total n = 7,003) but recorded at subsequent assessment waves, the value recorded subsequently was imputed into the ethnicity variable.

**Table S2.** *Descriptive statistics (Early life factors; EL)*

| | | N | Coding used |
|---|---|---|---|
| Comparative body size at age 10[a] | Thinner | 2,235 | 1 |
| | About average | 3,598 | 2 |
| | Plumper | 1,150 | 3 |
| | Missing | 20 | -- |
| Comparative height size at age 10[b] | Shorter | 1,315 | 1 |
| | About average | 3,687 | 2 |
| | Taller | 1,977 | 3 |
| | Missing | 24 | -- |
| Maternal smoke around the time of birth[c] | Yes | 1,986 | 0 |
| | No | 4,483 | 1 |
| | Missing | 534 | -- |
| Breastfed as a baby[d] | Yes | 4,409 | 1 |
| | No | 1,565 | 0 |
| | Missing | 1,029 | -- |
| | M(SE) | N | Range |
| Birth weight[e] | 3.38 (0.60) | 4,717 | .85 - 6 |

*Note.* Data coding accommodates consistent directions for the correlations among variables within the early life factors latent measurement model. Therefore, higher values are suggestive of favourable early life circumstances. All early life factors ([abcde]) where recorded at baseline. Where this information was missing, it was imputed from the subsequent assessments waves if recorded later (missing at baseline: n = 70[a], n = 72[b], n = 818[c], n = 1,378[d], n = 2,727[e] of total n = 7,003). All early life factors variables were kept with missing values, given that they were only evaluated in the full SEM model, where the ML estimator with missing values was used.

[ab]Morphometrics at age 10 recorded at the baseline (fields 1697, 1687). Participants responded to the question: When you were 10 years old, compared to average would you describe yourself as: Shorter / Taller / About average; When you were 10 years old, compared to average would you describe yourself as: Thinner / Plumper / About average.

[c]Maternal smoking was recorded at baseline (field 1787) for all participants, except those who indicated that they were adopted. Participants responded to the question: Did your mother smoke regularly around the time when you were born? Yes / No.

[d]Being breastfed as a bay was recorded at baseline (field 1677). Participants responded to the question: Were you breastfed when you were a baby? Yes / No.

[e]Birth weight (Kg) was recorded at baseline (field 20022). If birth-weight was known participants entered the weight.

**Adverse events in childhood and adulthood**

Childhood adversity (CA) and adult adversity (AA) were constructed as latent variables based on four (CA) and three (AA) variables selected from the respective questionnaires (Table 1). The full questionnaire for measuring CA uses the CTS-5 (Bellis et al., 2014; Bernstein et al., 1994; Glaesmer et al., 2013). This questionnaire is used in the clinical setting with diagnostics cut-offs (Glaesmer et al., 2013). Here, the questionnaire was used to determine a latent construct of CA using confirmatory factor analysis (CFA). This approach was considered more appropriate compared to a summative score because: (1) it allows allocation of varied factor loadings to each contributing variable, considering that different forms of child maltreatment or the number of experienced events may determine specific associations with mental health outcomes (Gilbert et al., 2009); (2) it accounts for both unique item variance and shared variance among items (which was expected with these items); (3) it provides a measure of goodness of fit. The full questionnaire measuring adult adversity included 5 bespoke questions, based on the national crime survey for being a victim of crime and adult domestic violence (Khalifeh et al., 2015). Here CFA was also used to determine a latent construct of adult adversity. For both latent factors, only the items that yielded correlation coefficients > .3 with at least one other variable were included in the analysis. In addition, item selection was also based on reasonable judgements related to the plausible association between variables.  For transparency, correlations with all CA / AA items are shown below (Table S3/4).

The CA questionnaire is based on the following items: “When I was growing up …”: (c1) “I felt loved”; (c2)”People in my family hit me so hard that it left me with bruises or marks”; (c3) “I felt that someone in my family hated me”; (c4) “Someone molested me (sexually)”; (c5) “There was someone to take me to the doctor if I needed it”. Similarly, the AA questionnaire is based on the following: “Since I was sixteen …”: (a1) “I have been in a confiding relationship”; (a2) “A partner or ex-partner deliberately hit me or used violence in any other way”; (a3) “Partner or ex-partner repeatedly belittled me to the extent that I felt worthless”; (a4) “Partner or ex-partner sexually interfered with me, or forced me to have sex against my wishes”; (a5) “There was money to pay the rent or mortgage when I needed it”.  All questions were rated on a Likert 5-point scale: never true, rarely true, sometimes true, often, very often true. Additionally, there was the option: “prefer not to answer”. This option was recoded as a missing value. The CA factor is based on (c1) (c2) (c3) and (c5). The AA factor is based on (a2) (a3) (a4). Figure S2.

**Table S3.** *Unadjusted Spearman correlations between items in the CA questionnaire on complete observations*

| **Childhood adversity** | | N††† | Item used (Y / N) | 1 | 2 | 3 | 4 | 5 |
|---|---|---|---|---|---|---|---|---|
| I felt that someone in my family hated me | EAc | 6948 | Y | -- | | | | |
| I felt loved a child†† | ENc | 6948 | Y | 0.38 | -- | | | |
| People in my family hit me so hard that it left me with bruises or marks | PAc | 6948 | Y | 0.38 | 0.29 | -- | | |
| Someone molested me (sexually) | SAc | 6948 | N | 0.16† | 0.13† | 0.13† | -- | |
| There was someone to take me to the doctor if I needed it†† | PNc | 6948 | Y | 0.19 | 0.34 | 0.16 | 0.08† | -- |

**Table S4.** *Unadjusted Spearman correlations between items in the AA questionnaire on complete observations*

| **Adult adversity** | | N††† | Item used (Y / N) | 1 | 2 | 3 | 4 | 5 |
|---|---|---|---|---|---|---|---|---|
| Partner or ex-partner repeatedly belittled me to the extent that I felt worthless | EAa | 6839 | Y | -- | | | | |
| I have been in a confiding relationship†† | ESa | 6839 | N | 0.14† | -- | | | |
| A partner or ex-partner deliberately hit me or used violence in any other way | PAa | 6839 | Y | 0.47 | 0.09† | -- | | |
| Partner or ex-partner sexually interfered with me, or forced me to have sex against my wishes | SAa | 6839 | Y | 0.35 | 0.09† | 0.33 | -- | |
| There was money to pay the rent or mortgage when I needed it†† | FHa | 6839 | N | 0.13† | 0.19† | 0.14† | 0.11† | -- |

*Note.* Tables S3/4: All correlation were significant at $p < .001$; †Correlation coefficients for items not included in the CFA. ††Items were reverse-coded. †††Correlation performed list-wise (further missing values in the excluded variables). ESa = (lack of) Emotional Support as an adult; FHa = financial hardship as an adult; see main manuscript for acronyms of used items

*Measurement models of adverse events*

Confirmatory factor analyses (CFA) were conducted to explore latent models of CA, AA and EL using the Satorra-Bntler robust estimator (Tables S5-S8). These models were evaluated separately before building the full SEM models. In addition the predicted factors were used in initial exploratory regressions (Table S9). The EL latent factor was constructed including all relevant variables irrespective of their correlation coefficient as these were not part of a validated scale. To improve model fit, the modification indices option available in STATA was used to add residual covariance paths. Dropping the residual covariances did not significantly alter the results. Coefficient loadings, fit statistics and covariance paths are shown below.

**Table S5**

| CA | Coefficient (standardized) | p-value | 95% CI | N |
|---|---|---|---|---|
| EAc | 0.76 | $p < .001$ | 0.72 – 0.79 | 7,003 |
| ENc | 0.58 | $p < .001$ | 0.55 – 0.61 | 7,003 |
| PAc | 0.62 | $p < .001$ | 0.58 – 0.65 | 7,003 |
| PNc | 0.29 | $p < .001$ | 0.24 – 0.33 | 7,003 |

**Table S6**

| AA | Coefficient (standardized) | p-value | 95% CI | N |
|---|---|---|---|---|
| EAa | 0.75 | $p < .001$ | 0.71 – 0.79 | 7,003 |
| PAa | 0.72 | $p < .001$ | 0.68 – 0.77 | 7,003 |
| SAa | 0.58 | $p < .001$ | 0.53 – 0.62 | 7,003 |

**Table S7**

| EL | Coefficient (standardized) | p-value | 95% CI | N |
|---|---|---|---|---|
| Body size age 10 | 0.17 | $p < .001$ | 0.11 – 0.23 | 4,144 |
| Height size age 10 | 0.25 | $p < .001$ | 0.18 – 0.33 | 4,144 |
| Maternal smoking | 0.06 | $p = .008$ | 0.02 – 0.11 | 4,144 |
| Breastfed as a baby | 0.10 | $p < .001$ | 0.05 – 0.15 | 4,144 |
| Birthweight | 0.65 | $p < .001$ | 0.46 – 0.83 | 4,144 |

**Table S8**

| Measurement model | $\chi^2$ | df | p-value | CFI | TLI | RMSEA | SRMR |
|---|---|---|---|---|---|---|---|
| CA | 2.72 | 1 | $p = .099$ | 0.99 | 0.99 | 0.02 | 0.01 |
| EL | 10.45 | 4 | $p = .033$ | 0.97 | 0.93 | 0.02 | 0.01 |

*Note.* Goodness of fit statistics could not be computed for the adult adversity model (just-identified model). Model fit was improved by allowing the following errors to correlate in the respective model: ENc and PNc; Maternal smoking and Breastfed as a baby.

**Figure S2**

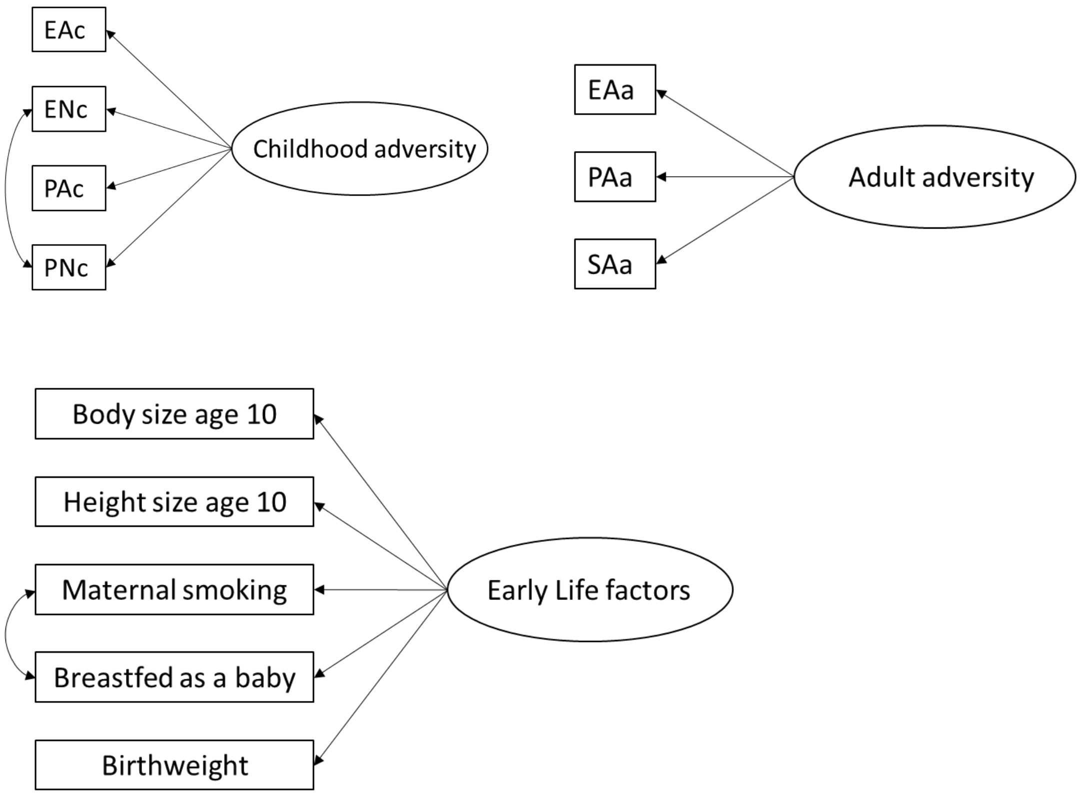
EAc
ENc
PAc
PNc
Childhood adversity
EAa
PAa
SAa
Adult adversity
Body size age 10
Height size age 10
Maternal smoking
Breastfed as a baby
Birthweight
Early Life factors

**Latent factors of white matter microstructure**

**Table S9.** *Unadjusted Pearson correlations between white matter microstructural measures (FA)*

| | 1 | 2 | 3 | 4 | 5 | 6 | 7 | 8 | 9 | 10 | 11 | 12 | 13 | 14 | 15 |
|---|---|---|---|---|---|---|---|---|---|---|---|---|---|---|---|
| Acoustic radiation (FA) | 1 | | | | | | | | | | | | | | |
| Anterior thalamic radiation (FA) | 0.843 | 1 | | | | | | | | | | | | | |
| Cingulate gyrus part of cingulum (FA) | 0.743 | 0.782 | 1 | | | | | | | | | | | | |
| Parahippocampal part of cingulum (FA) | 0.567 | 0.580 | 0.533 | 1 | | | | | | | | | | | |
| Corticospinal tract (FA) | 0.827 | 0.836 | 0.709 | 0.570 | 1 | | | | | | | | | | |
| Forceps major (FA) | 0.808 | 0.828 | 0.699 | 0.534 | 0.795 | 1 | | | | | | | | | |
| Forceps minor (FA) | 0.842 | 0.922 | 0.796 | 0.593 | 0.822 | 0.822 | 1 | | | | | | | | |
| Inferior fronto-occipital fasciculus (FA) | 0.871 | 0.931 | 0.788 | 0.608 | 0.840 | 0.858 | 0.928 | 1 | | | | | | | |
| Inferior longitudinal fasciculus (FA) | 0.891 | 0.906 | 0.778 | 0.611 | 0.854 | 0.870 | 0.909 | 0.969 | 1 | | | | | | |
| Middle cerebellar peduncle (FA) | 0.668 | 0.708 | 0.613 | 0.534 | 0.723 | 0.683 | 0.702 | 0.724 | 0.729 | 1 | | | | | |
| Medial lemniscus (FA) | 0.685 | 0.716 | 0.640 | 0.562 | 0.779 | 0.666 | 0.725 | 0.721 | 0.734 | 0.621 | 1 | | | | |
| Posterior thalamic radiation (FA) | 0.823 | 0.874 | 0.741 | 0.621 | 0.806 | 0.835 | 0.876 | 0.927 | 0.941 | 0.725 | 0.757 | 1 | | | |
| Superior longitudinal fasciculus (FA) | 0.876 | 0.899 | 0.779 | 0.585 | 0.840 | 0.832 | 0.902 | 0.923 | 0.929 | 0.693 | 0.715 | 0.876 | 1 | | |
| Superior thalamic radiation (FA) | 0.840 | 0.864 | 0.733 | 0.593 | 0.895 | 0.772 | 0.854 | 0.860 | 0.869 | 0.697 | 0.761 | 0.851 | 0.875 | 1 | |
| Uncinate fasciculus (FA) | 0.821 | 0.855 | 0.733 | 0.576 | 0.776 | 0.784 | 0.849 | 0.877 | 0.867 | 0.662 | 0.648 | 0.796 | 0.846 | 0.785 | 1 |

*Note.* A correlation matrix was generated for each type of tract measurement before constructing the latent factors. The strong correlations between white matter microstructural measures of fractional anisotropy are summarized in this table (IDPs averaged between hemispheres and saved as standardized residuals after controlling for confounds). All correlations were significant at $p < .001$.

**Table S10**. *Unadjusted Pearson correlations between white matter microstructural measures (MD)*

| | **1** | **2** | **3** | **4** | **5** | **6** | **7** | **8** | **9** | **10** | **11** | **12** | **13** | **14** | **15** |
|---|---|---|---|---|---|---|---|---|---|---|---|---|---|---|---|
| Acoustic radiation (MD) | 1 | | | | | | | | | | | | | | |
| Anterior thalamic radiation (MD) | 0.864 | 1 | | | | | | | | | | | | | |
| Cingulate gyrus part of cingulum (MD) | 0.860 | 0.918 | 1 | | | | | | | | | | | | |
| Parahippocampal part of cingulum (MD) | 0.733 | 0.774 | 0.744 | 1 | | | | | | | | | | | |
| Corticospinal tract (MD) | 0.865 | 0.914 | 0.911 | 0.774 | 1 | | | | | | | | | | |
| Forceps major (MD) | 0.728 | 0.757 | 0.718 | 0.670 | 0.758 | 1 | | | | | | | | | |
| Forceps minor (MD) | 0.842 | 0.908 | 0.925 | 0.724 | 0.873 | 0.689 | 1 | | | | | | | | |
| Inferior fronto-occipital fasciculus (MD) | 0.890 | 0.945 | 0.916 | 0.791 | 0.910 | 0.808 | 0.919 | 1 | | | | | | | |
| Inferior longitudinal fasciculus (MD) | 0.894 | 0.926 | 0.893 | 0.797 | 0.906 | 0.815 | 0.887 | 0.982 | 1 | | | | | | |
| Middle cerebellar peduncle (MD) | 0.584 | 0.631 | 0.578 | 0.591 | 0.621 | 0.532 | 0.583 | 0.635 | 0.639 | 1 | | | | | |
| Medial lemniscus (MD) | 0.778 | 0.791 | 0.800 | 0.723 | 0.846 | 0.658 | 0.799 | 0.811 | 0.806 | 0.592 | 1 | | | | |
| Posterior thalamic radiation (MD) | 0.827 | 0.878 | 0.851 | 0.742 | 0.870 | 0.754 | 0.865 | 0.920 | 0.919 | 0.616 | 0.822 | 1 | | | |
| Superior longitudinal fasciculus (MD) | 0.876 | 0.936 | 0.937 | 0.758 | 0.911 | 0.743 | 0.930 | 0.956 | 0.947 | 0.613 | 0.810 | 0.902 | 1 | | |
| Superior thalamic radiation (MD) | 0.858 | 0.933 | 0.945 | 0.746 | 0.931 | 0.710 | 0.921 | 0.912 | 0.894 | 0.609 | 0.835 | 0.896 | 0.949 | 1 | |
| Uncinate fasciculus (MD) | 0.867 | 0.911 | 0.862 | 0.797 | 0.881 | 0.784 | 0.841 | 0.930 | 0.937 | 0.630 | 0.761 | 0.819 | 0.884 | 0.839 | 1 |

*Note.* A correlation matrix was generated for each type of tract measurement before constructing the latent factors. The strong correlations between white matter microstructural measures of mean diffusivity are summarized in this table (IDPs averaged between hemispheres and saved as standardized residuals after controlling for confounds). All correlations were significant at $p < .001$.

**Table S11.** *LASSO feature selection and associated ß coefficients for IDPs identified by minimum one adversity variable.*

| | | EAc | ENc | PAc | PNc | EAa | PAa | SAa |
|---|---|---|---|---|---|---|---|---|
| **Frontal Lobe** | Inferior Frontal Gyrus, pars opercularis | | | | | | 0.006 | 0.008 |
| | Frontal Medial Cortex | | -0.008 | | | | | |
| | Middle Frontal Gyrus | | -0.011 | | | | | -0.0005 |
| | Central Opercular Cortex | | -0.011 | | | | | |
| | Frontal Operculum Cortex | | -0.001 | | | | | |
| | Superior Frontal Gyrus | | 0.006 | | | | | |
| | Supplementary Motor Cortex | | -0.016 | | | | -0.001 | |
| **Temporal lobe** | Middle Temporal Gyrus, anterior | | | | | | -0.003 | -0.002 |
| | Middle Temporal Gyrus, temporooccipital | | -0.006 | | | | | -0.007 |
| | Planum polare | | 0.008 | | | | | |
| | Temporal Pole | | 0.010 | | | | | |
| | Temporal Fusiform Cortex, posterior | | -0.001 | | | | | 0.006 |
| | Temporal Occipital Fusiform Cortex | | 0.017 | 0.006 | | | | |
| | Heschls Gyrus | | 0.013 | | | | 0.002 | |
| | Inferior Temporal Gyrus, anterior | | 0.004 | | | | | |
| | Inferior Temporal Gyrus, posterior | | | | | | 0.009 | |
| **Parietal Lobe** | Precuneous Cortex | | 0.015 | 0.008 | | | | 0.006 |
| | Postcentral Gyrus | | -0.020 | -0.008 | | | | |
| | Angular Gyrus | | 0.016 | 0.007 | | | -0.001 | |
| | Supramarginal Gyrus, anterior | | -0.009 | | | | | |
| | Supramarginal Gyrus, posterior | | -0.012 | | | | | |
| | Parietal Operculum Cortex | | | | | | | -0.002 |
| | Parietal Lobule, superior | | -0.003 | | | | | |
| **Occipital Lobe** | Occipital Pole | | 0.010 | | | 0.004 | 0.000 | |
| | Lingual Gyrus | | -0.017 | | | | | 0.001 |
| | Lateral Occipital Cortex, inferior | | | -0.003 | | | 0.003 | |
| | Lateral Occipital Cortex, superior | | 0.001 | | | | | |
| | Intracalcarine Cortex | | -0.017 | | | | | |
| | Supracalcarine Cortex | | -0.0001 | | | | | |
| **Cerebellum** | Cerebellum Crus I | | -0.002 | -0.009 | | | 0.000 | |
| | Cerebellum Crus II | | | | | | 0.003 | |
| | Cerebellum Lobules I-IV | | -0.010 | | | | | |
| | Cerebellum Lobule IX | | | | | -0.013 | | |
| | Cerebellum Lobule V | | | | | 0.017 | | 0.004 |
| | Vermis Crus I | | | | | 0.005 | 0.008 | 0.004 |
| | Vermis Crus II | | | | | | 0.002 | |
| | Vermis Lobule VI | | 0.019 | | | | | |
| | Vermis Lobule VIIIa | | | | | | | 0.002 |
| | Vermis Lobule VIIIb | | -0.004 | | | | | |
| | Vermis Lobule X | | | | | -0.028 | | -0.011 |
| | Cerebellum Lobule VIIIa | | -0.008 | | | | | |
| | Cerebellum Lobule X | | | -0.001 | | | | |
| **Basal ganglia** | Putamen | | | | | | 0.007 | 0.001 |
| | Pallidum | | 0.009 | | | -0.002 | -0.008 | -0.003 |
| | Ventral Striatum | -0.004 | 0.020 | | | | | |
| | Caudate | | 0.003 | | | 0.002 | | |
| **Subcortical / Other** | Paracingulate Gyrus | | 0.011 | | | | | |
| | Subcallosal Cortex | -0.006 | -0.032 | | | -0.005 | -0.008 | -0.005 |
| | Thalamus | | | | | 0.000 | 0.001 | 0.002 |
| | Amygdala | | | | | | | -0.001 |
| | Hippocampus | | -0.003 | | | | | 0.0001 |
| | Insular Cortex | | 0.006 | | | | | -0.001 |

*Note*. Shaded columns show results for CA. Under CA, features were primarily selected by the ENc variable, while PNc did not identify any non-redundant features.

**Table S12.** *Latent factors of grey matter structure - Unadjusted Pearson correlations*

*Unadjusted Pearson correlations between cerebellar grey matter structures*

| | **1** | **2** | **3** | **4** | **5** | **6** | **7** | **8** | **9** | **10** | **11** | **12** |
|---|---|---|---|---|---|---|---|---|---|---|---|---|
| Cerebellum Crus I | 1 | | | | | | | | | | | |
| Cerebellum Crus II | 0.413 | 1 | | | | | | | | | | |
| Cerebellum Lobules I-IV | 0.311 | 0.388 | 1 | | | | | | | | | |
| Cerebellum Lobule IX | 0.353 | 0.417 | 0.459 | 1 | | | | | | | | |
| Cerebellum Lobule V | 0.365 | 0.388 | 0.766 | 0.502 | 1 | | | | | | | |
| Vermis Crus II | 0.272 | 0.484 | 0.326 | 0.278 | 0.363 | 1 | | | | | | |
| Vermis Lobule VI | 0.323 | 0.311 | 0.401 | 0.352 | 0.487 | 0.528 | 1 | | | | | |
| Vermis Lobule VIIIa | 0.352 | 0.492 | 0.454 | 0.608 | 0.517 | 0.549 | 0.459 | 1 | | | | |
| Vermis Lobule VIIIb | 0.321 | 0.449 | 0.428 | 0.711 | 0.453 | 0.381 | 0.377 | 0.777 | 1 | | | |
| Vermis Lobule X | 0.276 | 0.337 | 0.415 | 0.528 | 0.375 | 0.261 | 0.266 | 0.409 | 0.431 | 1 | | |
| Cerebellum Lobule VIIIa | 0.315 | 0.624 | 0.474 | 0.700 | 0.528 | 0.363 | 0.403 | 0.671 | 0.624 | 0.445 | 1 | |
| Cerebellum Lobule X | 0.281 | 0.248 | 0.264 | 0.172 | 0.287 | 0.194 | 0.241 | 0.208 | 0.219 | 0.104 | 0.235 | 1 |

*Unadjusted Pearson correlations between basal ganglia structures*

| | **1** | **2** | **3** | **4** |
|---|---|---|---|---|
| Pallidum (GMV) | 1 | | | |
| Ventral striatum (GMV) | 0.140 | 1 | | |
| Putamen (GMV) | 0.260 | 0.612 | 1 | |
| Caudate (GMV) | 0.214 | 0.322 | 0.464 | 1 |

*Unadjusted Pearson correlations between frontal lobe structures*

| | **1** | **2** | **3** | **4** | **5** | **6** | **7** |
|---|---|---|---|---|---|---|---|
| Inferior Frontal Gyrus, pars opercularis | 1 | | | | | | |
| Frontal Medial Cortex | 0.125 | 1 | | | | | |
| Middle Frontal Gyrus | 0.155 | 0.142 | 1 | | | | |
| Central Opercular Cortex | 0.280 | 0.170 | 0.227 | 1 | | | |
| Frontal Operculum Cortex | 0.376 | 0.152 | 0.140 | 0.206 | 1 | | |
| Superior Frontal Gyrus | 0.161 | 0.134 | 0.091 | 0.245 | 0.231 | 1 | |
| Supplementary Motor Cortex | 0.102 | 0.116 | 0.124 | 0.152 | 0.138 | 0.291 | 1 |

*Unadjusted Pearson correlations between temporal lobe structures*

| | **1** | **2** | **3** | **4** | **5** | **6** |
|---|---|---|---|---|---|---|
| Planum polare | 1 | | | | | |
| Temporal Pole | 0.140 | 1 | | | | |
| Temporal Fusiform Cortex, posterior | 0.297 | 0.161 | 1 | | | |
| Heschls Gyrus | 0.327 | 0.047† | 0.200 | 1 | | |
| Inferior Temporal Gyrus, anterior | 0.112 | 0.295 | 0.168 | 0.164 | 1 | |
| Inferior Temporal Gyrus, posterior | 0.186 | 0.187 | 0.532 | 0.211 | 0.358 | 1 |

†$p < .01$

*Unadjusted Pearson correlations between parietal lobe structures*

| | 1 | 2 | 3 | 4 | 5 | 6 | 7 |
|---|---|---|---|---|---|---|---|
| Precuneous Cortex | 1 | | | | | | |
| Postcentral Gyrus | 0.189 | 1 | | | | | |
| Angular Gyrus | 0.224 | 0.188 | 1 | | | | |
| Supramarginal Gyrus, anterior | 0.173 | 0.408 | 0.169 | 1 | | | |
| Supramarginal Gyrus, posterior | 0.185 | 0.212 | 0.451 | 0.481 | 1 | | |
| Parietal Operculum Cortex | 0.237 | 0.135 | 0.067 | 0.280 | 0.152 | 1 | |
| Parietal Lobule, superior | 0.224 | 0.201 | 0.148 | 0.063 | 0.191 | 0.083 | 1 |

*Unadjusted Pearson correlations between occipital lobe structures*

| | 1 | 2 | 3 | 4 | 5 | 6 |
|---|---|---|---|---|---|---|
| Occipital Pole | 1 | | | | | |
| Lingual Gyrus | 0.282 | 1 | | | | |
| Lateral Occipital Cortex, inferior | 0.276 | 0.187 | 1 | | | |
| Lateral Occipital Cortex, superior | 0.190 | 0.152 | 0.198 | 1 | | |
| Intracalcarine Cortex | 0.520 | 0.398 | 0.267 | 0.046† | 1 | |
| Supracalcarine Cortex | 0.249 | 0.155 | 0.241 | 0.062 | 0.408 | 1 |

†$p < .01$

*Note.* A correlation matrix was generated for each grey matter category before constructing the latent factors. The correlations among grey matter IDPs within each category is summarized in these tables. All IDPs represent standardized residuals after controlling for confounds. Unless flagged (†), all correlations were significant at p < .001.

**Table S13.** *Model fit table for 20 structural models, each including a different brain measure*

| Brain measure | $X^2$ | df | p-value | CFI | TLI | RMSEA | SRMR |
|---|---|---|---|---|---|---|---|
| Total GM | 165.84 | 59 | $p < .001$ | 0.98 | 0.97 | 0.02 | 0.03 |
| Total WM | 152.58 | 59 | $p < .001$ | 0.98 | 0.97 | 0.02 | 0.02 |
| gFL | 140.56 | 59 | $p < .001$ | 0.98 | 0.98 | 0.02 | 0.02 |
| gSTG | 150.65 | 59 | $p < .001$ | 0.98 | 0.97 | 0.02 | 0.02 |
| gITG | 139.45 | 59 | $p < .001$ | 0.98 | 0.98 | 0.02 | 0.02 |
| gPL | 153.04 | 59 | $p < .001$ | 0.98 | 0.97 | 0.02 | 0.02 |
| gOL | 147.62 | 59 | $p < .001$ | 0.98 | 0.97 | 0.02 | 0.02 |
| gCBL | 161.75 | 59 | $p < .001$ | 0.98 | 0.97 | 0.02 | 0.02 |
| gBG | 153.90 | 59 | $p < .001$ | 0.98 | 0.97 | 0.02 | 0.02 |
| gFA | 140.52 | 59 | $p < .001$ | 0.98 | 0.98 | 0.02 | 0.02 |
| gMD | 146.71 | 59 | $p < .001$ | 0.98 | 0.97 | 0.02 | 0.02 |
| Paracingulate Gyrus | 140.65 | 59 | $p < .001$ | 0.98 | 0.98 | 0.02 | 0.02 |
| Subcallosal Cortex | 140.90 | 59 | $p < .001$ | 0.98 | 0.98 | 0.02 | 0.02 |
| Thalamus | 153.97 | 59 | $p < .001$ | 0.98 | 0.97 | 0.02 | 0.02 |
| Amygdala | 150.80 | 59 | $p < .001$ | 0.98 | 0.97 | 0.02 | 0.02 |
| Hippocampus | 153.47 | 59 | $p < .001$ | 0.98 | 0.97 | 0.02 | 0.02 |
| Insular Cortex | 161.59 | 59 | $p < .001$ | 0.98 | 0.97 | 0.02 | 0.02 |
| Vermis Crus I | 145.69 | 59 | $p < .001$ | 0.98 | 0.97 | 0.02 | 0.02 |
| Middle Temporal Gyrus, anterior | 155.07 | 59 | $p < .001$ | 0.98 | 0.97 | 0.02 | 0.02 |
| Middle Temporal Gyrus, temp-occ. | 143.29 | 59 | $p < .001$ | 0.98 | 0.97 | 0.02 | 0.02 |
| Temp-occ. Fusiform Cortex | 148.87 | 59 | $p < .001$ | 0.98 | 0.97 | 0.02 | 0.02 |

*Note.* All structural models showed good fit of the data. The first column indicates the brain measure used in each of the 20 structural equation models. The Satorra-Bentler estimator was used for all models, and therefore the fit statistics above were adjusted. Model fit was improved by adding additional covariance paths (Figure S2). GM = grey matter; WM = white matter; gFL = global frontal lobe; gSTG = global superior temporal gyrus; gITG = global inferior temporal gyrus; gPL = global parietal lobe; gOL = global occipital lobe; gCBL = global cerebellum; gBG = global basal ganglia; gFA = global fractional anisotropy; gMD = global mean diffusivity.
